# A session-aware test protocol, energy-based strand pairing and a measured crossing-resolution limit for automated DNA knot identification in atomic force microscopy images

**DOI:** 10.64898/2026.11.12.766324

**Authors:** Musabbir Hasan Sammak, Khalid Been Md Badruzzaman Biplob, Tapushe Rabaya Toma, Imran Mahmud

## Abstract

Atomic force microscopy (AFM) resolves the crossings of single knotted and linked DNA molecules, so the knot type can in principle be read from one image. TopoStats, to our knowledge the only automated pipeline for this task, assigns the correct type to fewer than 8% of the knots and catenanes in a public dataset of 721 molecules imaged in 20 sessions. Image classifiers appear to do better, but 18 of the 20 sessions contain a single knot type, so a classifier can score well by recognizing the session. We present three tools for evaluating and tracing these molecules. First, a session-aware protocol holds out whole imaging sessions, scores only the four classes not separable by size, and uses a probe to test whether images identify their session. Three classifiers reached 58–67% balanced accuracy on knots on their training sessions but only 27–32% under this protocol, near the 25% chance level, and the probe identified the session above chance in all eight tests. Second, an energy-based method connects strands through crossings by DNA length and stiffness. It traced the loops of 84% of knots and catenanes correctly, compared with 32% for TopoStats. Third, we measured the crossing-resolution limit of the pipeline. Labels derived from the known handedness of the knots gave the correct upper strand at 173 crossings without manual annotation, and the existing width rule ordered 137 of them (79%) correctly. A convolutional network did not exceed it. With the new pairing, the knot type was still correct for only 6–8% of knots and catenanes, because 72% of those with a valid pairing had lost crossings when closely spaced crossings were merged. Classifiers for AFM images should therefore be tested on new sessions, and further progress depends on resolving dense crossings. The protocol, labels and code are openly available.

## Introduction

Many DNA molecules are circular, with the two ends of the double helix joined into a closed loop. Examples include bacterial plasmids, human mitochondrial DNA [1] and extrachromosomal circular DNA, which separates from the chromosomes and occurs both in healthy tissue [2] and in nearly half of human cancers [3]. A closed loop of DNA can become entangled. A loop that is entangled with itself and cannot be pulled into a simple ring without cutting the DNA is called a knot. Two or more loops that pass through each other, like the links of a chain, form a catenane. Knots and catenanes come in different types, depending on how the DNA is entangled. These types are informative. Each enzyme that cuts and rejoins DNA leaves behind knots or catenanes of particular types, so the types found after a reaction show how the enzyme binds, cuts and rejoins the DNA [4,5]. Knotting also affects how DNA functions in the cell. For example, the enzyme T7 RNA polymerase cannot transcribe highly knotted DNA [6], and knotting in bacteria impairs plasmid replication and increases mutation rates [7]. Determining the knot type of individual molecules is therefore valuable for studying DNA replication, recombination and the enzymes involved [8].

In this paper, the term knot type is used for the type of both knots and catenanes. Knot types are named by their crossings. In a drawing of a knot, a crossing is a point where the DNA passes over itself. The same knot can be drawn in many ways, and the smallest number of crossings that any drawing of it can have is its minimal crossing number. In the standard notation [9], the main number of a knot type gives this minimal crossing number. A subscript distinguishes types that have the same number, so 5₁ and 5₂ are two different knots with five crossings. For a catenane, a superscript gives the number of loops. This study concerns the five types shown in Fig 1. These are the unknot 0₁, a simple ring without crossings, the trefoil 3₁, the torus knot 5₁, the twist knot 5₂ and the catenane 4²₁, whose two loops are linked by four crossings.

**Fig 1.**
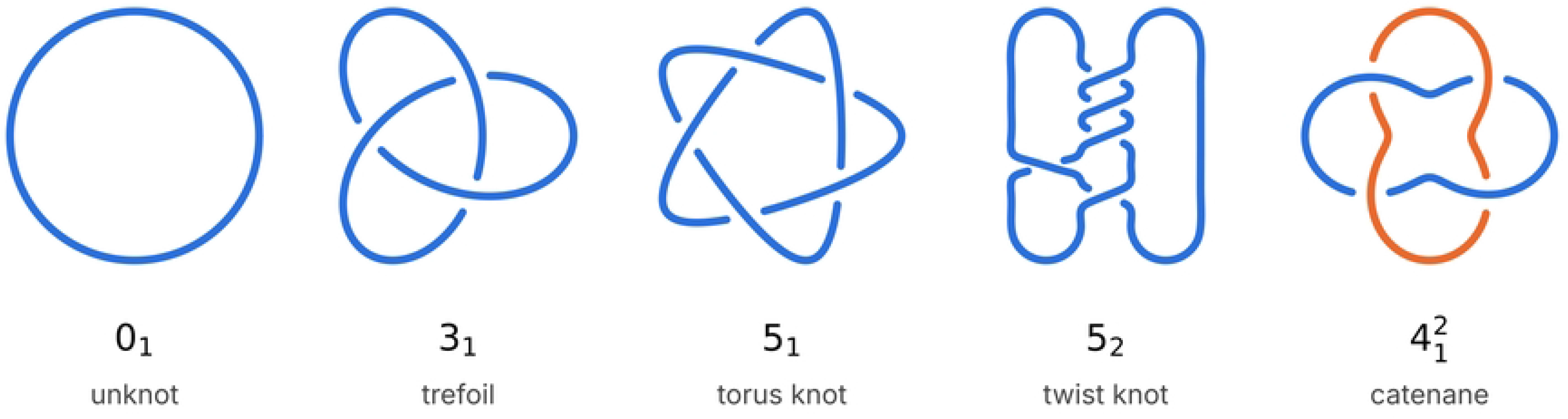
The five knot types studied. Each type is drawn as a minimal diagram, and a gap marks the lower strand at each crossing. The diagrams were generated from closed three-dimensional curves, and Topoly confirmed the type of each curve. The crossing signs match the chirality of the samples, positive for 3₁, 5₁ and 4²₁ and negative for 5₂. The two loops of the catenane are shown in different colors. The code that draws and checks the diagrams was written with the assistance of Claude (Anthropic).

Knot types have traditionally been identified by gel electrophoresis or by electron microscopy. Gel electrophoresis measures a whole population of molecules rather than individual ones, and electron microscopy requires the DNA to be coated and shadowed with metal [8]. High-resolution atomic force microscopy (AFM) in liquid avoids both limits. It images single, uncoated molecules and resolves the individual crossings of plasmid-sized DNA, so the knot type can in principle be read from a single image [8,10]. Reading the knot type by hand, however, is slow and subject to observer bias, and this has led to efforts to automate it. Several tools trace DNA molecules in AFM images automatically [11–13]. One of them, TopoStats, also follows the DNA through crossings [14], and Holmes et al. [8] extended it to identify the knot type. To our knowledge, this pipeline remains the only automated one that does so. At each crossing, the pipeline places on top the strand whose height profile across the crossing is wider, a decision we call the width rule. It then builds a three-dimensional model of the molecule. From this model, it identifies the knot type with the HOMFLY-PT polynomial [15], a mathematical invariant that distinguishes knot types, computed with the software Topoly [16].

Holmes et al. [8] released their images together with the TopoStats output for every traced molecule [17]. Each sample contains a single knot type of known identity, so the type that TopoStats assigns to each molecule can be checked against the true type. When broken and overlapping molecules are excluded, TopoStats assigns the correct type to only 44 of 588 knots and catenanes (7.5%). This accuracy is lower than the one Holmes et al. [8] reported, because they removed molecules with unresolvable crossings before classifying them. Even so, it shows that the fully automatic pipeline is not yet usable for these samples. It also raises the question of which step of the pipeline causes the errors.

Machine learning could improve this analysis. Neural networks identify knot types with high accuracy when the full three-dimensional shape of a simulated molecule is known [18], and recent work on AFM images has trained networks on simulated data [19,20]. An AFM image, however, shows only the height of a molecule seen from above, so learned methods must be tested on measured images. Three obstacles stand in the way of improving the pipeline, whether with machine learning or without it. The first obstacle concerns how AFM images are acquired. Images are collected in imaging sessions, each typically one day on the instrument, and sessions can differ in tip condition, surface preparation and image contrast. In the dataset used here, 18 of the 20 sessions contain molecules of only one knot type. The appearance of a session therefore predicts the knot type. When molecules are divided into training and test sets without regard to session, the same sessions appear in both sets. A classifier can then score well by recognizing the session rather than the knot. This is called shortcut learning [21], and we refer to this case as session leakage. It is a form of data leakage, a common cause of inflated performance in machine-learning-based science [22,23]. Comparable effects have been reported for chest radiographs, where models learned to identify the hospital of origin [24], and in ecology, where models must be tested on sites not used for training [25]. For AFM images of DNA topology, the size of this effect has not been measured.

The second obstacle concerns how strands are connected through crossings. In an AFM image, the DNA at a crossing appears as a small area where two strands overlap, which TopoStats marks as a crossing region. Before the pipeline can decide which strand lies on top, it must connect each strand that enters a crossing region to a strand that leaves it. TopoStats makes these connections from the directions of the strands alone. A wrong connection, however, changes the loops that the traced molecule forms. It can join two loops into one, split one loop into two, or produce loops of the wrong length. Every intact molecule of a sample contains the same length of DNA, so only the connections that give loops of the expected length can be correct. TopoStats does not use this constraint.

The third obstacle is the lack of labels for individual crossings. The step that decides which strand lies on top cannot be tested without the correct answer at each crossing, and such answers are not available at scale. Every molecule carries the knot type of its sample as a label, but the upper strand at each of its crossings is unknown. Labeling crossings by hand is slow, and the labels are uncertain where the two strands differ little in height. Holmes et al. [8] calibrated the width rule on only 83 hand-labeled crossings.

This paper presents three methods, one for each obstacle, and evaluates each on the public dataset of Holmes et al. [8]. The first method addresses session leakage. It is a session-aware test protocol with four parts. It holds out whole imaging sessions during testing, and it scores only the four knot types that have the same DNA length and therefore cannot be told apart by size. It also compares learned models with TopoStats on the same molecules, and it tests directly whether images contain features that identify their session. The second method addresses the connection of strands. This energy-based pairing scores every possible set of connections with an energy that penalizes loops of the wrong length and strands that bend sharply, and it keeps the set with the lowest energy. The third method addresses the missing crossing labels. Each knot sample contains only one of the knot’s two mirror images, and this mirror image determines which strand lies on top at every crossing. The correct upper strand is therefore known without labeling by hand. These labels are used to measure how often the width rule is correct. The third method also counts the crossings that are missed because closely spaced crossings merge into one region, which shows how strongly crossing resolution limits the pipeline. Together, these methods show that whole-molecule classifiers learn the imaging session rather than the knot type, and that energy-based pairing corrects most loop errors. They also show that the pipeline is limited mainly by the merging of closely spaced crossings rather than by errors in their order. The protocol, the crossing labels and all code are openly available.

## Materials and methods

### Dataset

This study used a public dataset that Holmes et al. [8] released and Gamill et al. [17] deposited. It contains 402 high-resolution AFM height images of seven DNA preparations of known knot type, together with the TopoStats output for every image. The knots were produced by Xer recombination as described in [8]. One plasmid gives twist knots, including the 5₂ knot, and a second plasmid gives the torus knots 3₁ and 5₁ by the same reaction. Each knot sample contains a single mirror form [8]. All knots and unknots are made from the same 2,260 bp plasmid, and the catenanes are right-handed 4²₁ links of a 1,253 bp circle and a 398 bp circle. The unknot controls are the same plasmid without knots. Both the unknot controls and the catenanes were imaged in two states, supercoiled, in which the closed DNA is twisted on itself, and nicked, in which one strand is cut so that this twisting is released (Table 1). The images were acquired in 2022 in a Ni²⁺ buffer and in 2023 in a Mg²⁺/Ni²⁺ buffer, with pixel sizes from 0.29 to 0.98 nm, so imaging sessions can differ in buffer and pixel size as well as in tip condition and surface preparation.

**Table 1.** Molecules and imaging sessions per class.

| Class | Preparations | Images | Molecules kept | Sessions | Molecules in largest session, n (%) | TopoStats correct type, n (%) |
|---|---|---|---|---|---|---|
| **0<sub>1</sub> (unknot)** | supercoiled and nicked plasmid | 81 | 133 | 3 | 67 (50) | 115 (86.5) |
| **3<sub>1</sub> (trefoil)** | pINV product | 39 | 41 | 2 | 38 (93) | 18 (43.9) |
| **5<sub>1</sub> (torus knot)** | pINV product | 43 | 63 | 4 | 42 (67) | 4 (6.3) |
| **5<sub>2</sub> (twist knot)** | pDIR product | 39 | 56 | 4 | 29 (52) | 5 (8.9) |
| **4<sup>2</sup><sub>1</sub> (catenane)** | supercoiled and nicked catenane | 200 | 428 | 9 | 98 (23) | 17 (4.0) |
| **Total** | 7 preparations | 402 | 721 | 20 |  | 159 (22.1) |

### Selection of molecules and definition of imaging sessions

TopoStats detected 759 molecules in the 402 images. The file name of each image contains its acquisition date, and all images from one date were treated as one imaging session, which gave 20 sessions. Session and knot type are strongly linked. Among the molecules kept for analysis, 18 of the 20 sessions contain molecules of only one knot type, and 38 of the 41 trefoils were imaged in a single session (Table 1).

TopoStats detects every object in an image, including broken molecules, molecules cut off by the image border and molecules lying on top of each other. Such objects were removed with a length filter and a border check. All intact molecules of one kind contain the same amount of DNA, so they have nearly the same traced length. For knots and unknots, the median traced length was 756–765 nm across preparations, and 758 nm was used as the expected length. For catenanes, the medians were 553–555 nm, and 554 nm was used. A molecule was kept if its traced length lay within 10% of either expected length and it did not touch the image border. The filter does not use the knot type of the sample, so it could also be applied to images of unknown topology. It left 721 molecules (Table 1), which were used in all analyses of whole molecules.

### Overview of the evaluated pipeline

Fig 2 shows the steps of the pipeline and how each was evaluated. TopoStats reduces each molecule to a skeleton, a line one pixel wide that runs along the DNA, and its crossing regions split the skeleton into segments. The segment ends at each region are then paired into strands, the upper strand at each crossing is decided, and the knot type is computed from both. A whole-molecule image classifier would bypass these steps. We evaluated such classifiers with the session-aware protocol, replaced the pairing step with the energy-based method, and measured how far crossing resolution limits the final knot type. In every comparison, the method under study was compared with TopoStats on the same molecules or crossings.

**Fig 2.**
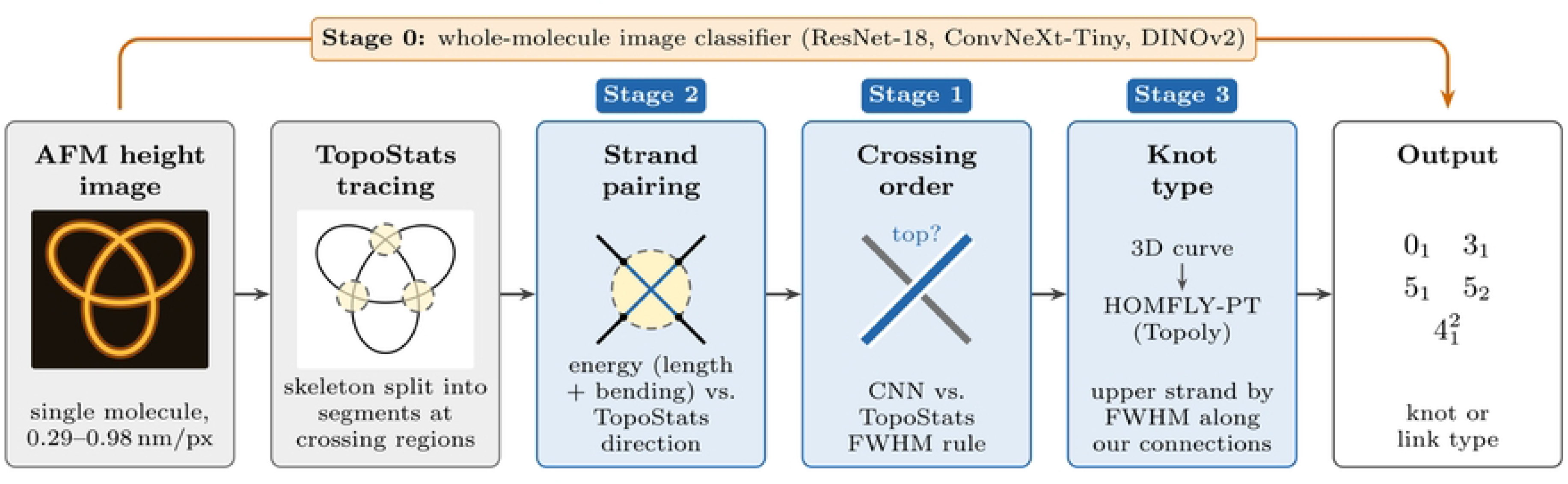
The analysis pipeline and the evaluated steps. Stage 2 pairs the segment ends at each crossing region into strands, Stage 1 decides the upper strand at each crossing, and Stage 3 computes the knot type. Stage 0 replaces these steps with a whole-molecule image classifier. The stages are numbered in the order in which they were evaluated. FWHM, full width at half maximum. CNN, convolutional neural network. The diagram was drawn with code written with the assistance of Claude (Anthropic).

### Session-aware test protocol

The session-aware test protocol has four parts, and each part prevents a specific way in which results on AFM images can be overstated. First, whole imaging sessions are held out during testing, so that a classifier cannot recognize a session it has seen in training. Second, performance is reported on the knot types that cannot be separated by size, so that an easy distinction cannot hide a failure on the others. Third, learned models are compared with TopoStats on the same molecules with a paired test. Fourth, a probe tests directly whether images identify their session.

#### Grouped cross-validation

All learned models were evaluated by five-fold cross-validation. The molecules were divided into five groups, called folds, and each fold was predicted by a model trained on the other four. Every molecule was therefore predicted exactly once, by a model that had not seen it. The folds were formed with StratifiedGroupKFold in scikit-learn [26]. This function keeps related molecules together in one fold and keeps the share of each knot type similar across folds. Two ways of grouping were compared. In the session split, all molecules from one imaging session were kept in the same fold, so every test fold contained only sessions that were absent from training. In the image split, only the molecules from one AFM image were kept together, so other images from the same session could appear in both training and test folds. The image split corresponds to the usual practice and was used only to measure how far it overestimates performance. The session folds are listed in S3 Table.

#### Knots-only metrics

The classifiers assign each molecule to one of five classes, the five knot types of Fig 1. Catenanes contain 554 nm of DNA and all other molecules 758 nm, so catenanes can be recognized from size alone. A high overall score could therefore hide a failure on the other four types. The primary metric was the knots-only balanced accuracy, computed on the 293 molecules of types 0₁, 3₁, 5₁ and 5₂. For each of these types, we measured the share of its molecules that were classified correctly, called its recall, and then averaged the four recalls.

These four types have the same DNA length and cannot be separated by size, so the chance level is 25%. The classifiers still chose among all five classes, and a knot classified as a catenane counted as an error, so the metric scores the same models that would be used in practice. As a second measure, we reported the knots-only macro AUC, which uses the predicted probabilities rather than only the predicted class. For each of the four types, the area under the receiver operating characteristic curve (AUC) measures how well the probabilities separate that type from the other three, and the four values were averaged. A value of 0.5 means that the probabilities carry no information about knot type. Accuracy, balanced accuracy and macro-F1 over all five classes were also reported.

#### Session probe

A low score on held-out sessions does not by itself prove session leakage, because it could also arise from too little training data. The session probe therefore tests directly whether images carry features that identify their session. A classifier was trained to predict the imaging session from images of a single preparation, for example only the nicked catenanes. All molecules of one preparation have the same knot type and the same supercoiling, so only the imaging conditions differ between their sessions. If the probe predicts the session above chance, the images must contain session features. The probe used the frozen DINOv2 features and the logistic regression described below, with five-fold cross-validation grouped by image. It was run for each preparation, and separately for each imaging buffer when a preparation was imaged in both buffers. Sessions with fewer than five images were left out, because such a session cannot appear in both training and test folds. Performance was measured as the balanced accuracy over sessions, whose chance level is one divided by the number of sessions. To obtain a *p* value, the session labels were shuffled between whole images 200 times, and the cross-validation was repeated after each shuffle.

#### Classifiers used to test the protocol

The protocol was applied to three whole-molecule classifiers of different design, a standard convolutional network, a more recent convolutional network and a self-supervised vision transformer. Three designs were used so that an effect shared by all three could be attributed to the data rather than to one architecture.

Each classifier received an image of a single molecule. The molecule was cut out of its AFM image with the mask produced by TopoStats. The mask was first enlarged by about 2 nm so that the edges of the DNA were kept, and all pixels outside it were set to zero. Every molecule was resampled to 1 nm per pixel and centered in an image covering 256 × 256 nm. Heights were limited to the range −1 to 5 nm and scaled linearly to between 0 and 1. Because the networks were pretrained on color photographs from ImageNet, each image was copied into three color channels, scaled with the ImageNet mean and standard deviation, and resized to 224 × 224 pixels.

ResNet-18 [27] and ConvNeXt-Tiny [28] were initialized with ImageNet weights, and all their layers were then trained on the molecule images in PyTorch [29]. Both were trained for 30 epochs with the AdamW optimizer (learning rate 3 × 10⁻⁴, weight decay 10⁻⁴), a cosine learning-rate schedule and batches of 32 images. The loss was cross-entropy, weighted by the inverse frequency of each class in the training folds so that rare classes counted as much as common ones. Training images were randomly rotated by 0–360° and shifted by up to 5%. Each test image was predicted at four rotations, by 0°, 90°, 180° and 270°, and the predictions were averaged. The third classifier used DINOv2 ViT-S/14 [30], in the version vit_small_patch14_dinov2.lvd142m from timm [31], without further training. Its features were averaged over the same four rotations and classified by class-balanced logistic regression (*C* = 1). All settings were fixed in advance and were not tuned on any fold, so that no test fold could influence the choice of model. Control analyses of flip augmentation, training length, input encoding and sampling density are described in S1 Appendix, and the results of all runs are shown in S1 Fig.

### Energy-based strand pairing

At each crossing region, the strands that enter must be connected to the strands that leave (Fig 3A). TopoStats pairs the branches of a region by their directions [8]. The energy-based method instead uses two physical properties of the sample. All intact molecules of one preparation contain the same length of DNA, and DNA is stiff on the scale of a crossing, so a strand tends to continue straight through it [32].

**Fig 3.**
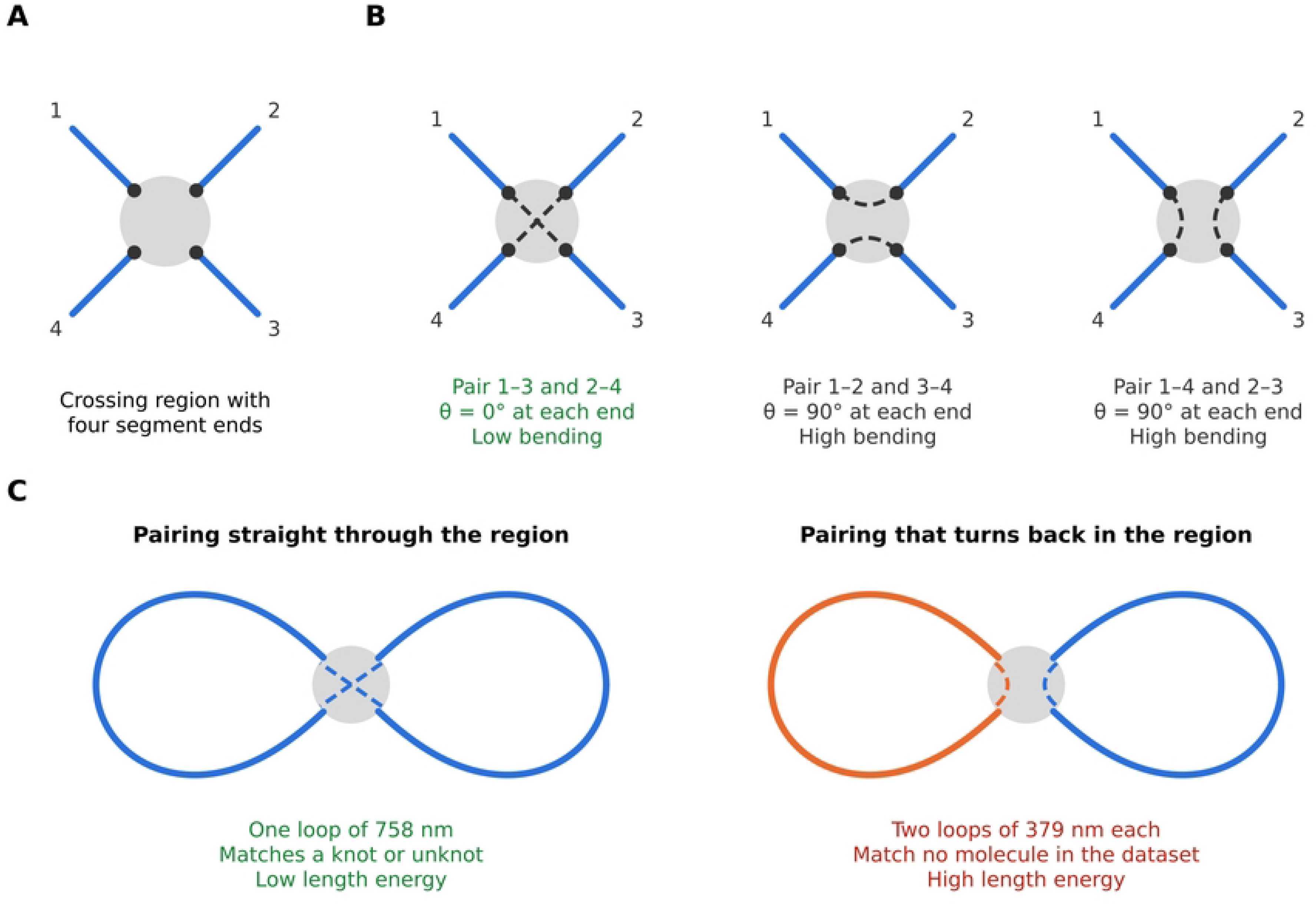
Energy-based strand pairing. (A) A crossing region where four segment ends meet. (B) The three possible pairings of these ends. Pairing straight through the region bends the strands least (*θ* = 0°), and pairings that turn back bend each strand by 90°. (C) The pairing also sets the loops of the whole molecule. Pairing straight through gives one loop of 758 nm, the length of a knot or unknot in the dataset. Pairing that turns back gives two loops of 379 nm, which match no molecule in the dataset. The method keeps the pairing with the lowest total energy (Eq 1). The diagram was drawn with code written with the assistance of Claude (Anthropic).

For each molecule, the TopoStats skeleton was split at the crossing regions into segments. Skeletonization can leave spurs, short branches of the skeleton that stick out from the edge of a crossing region and end a few pixels away. Spurs shorter than 5 pixels were ignored. Otherwise each spur would count as an extra strand end at its crossing region, and a region with an odd number of ends cannot be paired. The length of each segment was measured along its path after smoothing with a 7-pixel moving average, which removes the staircase bias of pixel paths. The direction of each segment end, which is needed to measure how sharply a strand bends through a crossing, was taken as the line from the end to the point 15 pixels further along the segment.

These three values were fixed before the pairing was applied to the data and were not tuned. At pixel sizes of 0.29–0.98 nm they correspond to about 1.5–5 nm, 2–7 nm and 4–15 nm. The spur threshold is shorter than the apparent width of DNA in the images, and the smoothing window and direction length are much shorter than the persistence length of DNA (about 50 nm) [32], over which DNA is nearly straight.

A pairing *P* joins the segment ends at every region in pairs (Fig 3B). Following the segments through these pairs gives a set of closed loops with lengths *ℓ*_1_ ≥ *ℓ*_2_ ≥ …, where each loop length includes the straight path through every region that the loop passes. Each pairing was scored by the energy

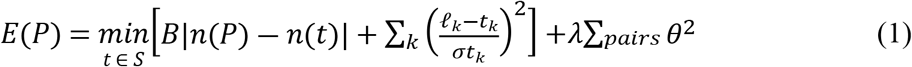

where *S* = {(758), (418, 136)} nm is the set of molecular species in the dataset, taken from the median loop lengths in the released TopoStats output, *n* is the number of loops, *B* = 1,000, *σ* = 0.05 and *λ* = 1. The angle *θ* measures how far a strand deviates from a straight path through a region, and the last sum runs over all joined pairs. The length term requires the loops to have the lengths of a real molecule (Fig 3C), and the large value of *B* makes a wrong number of loops much more costly than any length deviation. The bending term favors strands that continue straight through a crossing (Fig 3B).

The method is not told which sample a molecule comes from. For each pairing, it computes the length term twice, once assuming the molecule is a single 758 nm circle (a knot or unknot) and once assuming it is a 418 nm and a 136 nm circle (a catenane), and it keeps the lower value. The same method can therefore trace knots and catenanes without being told which is which. A region with four segment ends can be paired in three ways and a region with six ends in fifteen, so a molecule with several regions has many possible pairings. All of them were scored, and the pairing with the lowest energy was kept.

For the few molecules with more than 200,000 possible pairings, only the five pairings with the least bending were kept at each region, and only their combinations were scored, so that the search finished in reasonable time. No valid pairing exists when the skeleton breaks off without reaching a crossing region, when a piece of skeleton is not connected to any crossing region, or when a crossing region has an odd number of segment ends, because the ends can then not all be joined in pairs.

To judge a pairing, we counted the loops that it traced for each molecule and measured their lengths. For energy-based pairing, these are the loops formed by following the segments through the chosen connections. For TopoStats, they are the loops stored in its output file. A knot or unknot was counted as correct when it was traced as one loop of 682–834 nm, which is within 10% of 758 nm. A catenane was counted as correct when it was traced as two loops, the larger of 376–460 nm, which is within 10% of 418 nm, and the smaller of 110–165 nm. The loops traced by TopoStats were judged by the same rule. This test favors the energy-based method, because the method was built to produce loops of exactly these lengths. We therefore also judged the pairing by the knot type it leads to. That test is fair to both methods, because the energy contains no information about knot type. Both tests used the 721 molecules of Table 1, and a molecule without a valid pairing was counted as incorrect.

### Measuring the crossing-resolution limit

#### Crossing labels from knot handedness

To test how often the width rule places the correct strand on top, the correct upper strand must be known at each crossing. We obtained it from the known handedness of the knot samples, without labeling by hand. Deriving labels from such domain knowledge is known as weak supervision [33]. The method uses the sign of a crossing. Once a direction of travel is chosen along the DNA, every crossing has a sign, positive or negative, that depends on which strand lies on top.

Exchanging the upper and lower strands reverses the sign, so the mirror image of a knot has all its signs reversed. When a 3₁, 5₁ or 5₂ knot is drawn with its minimal number of crossings, all its crossings have the same sign. Which of the two mirror forms a knot takes, called its handedness or chirality, decides whether this sign is positive or negative.

Each sample contains a single mirror form, so the expected sign of its crossings can be known. Holmes et al. [8] reported that all crossings of the 5₁ knots are positive and that all crossings of the 5₂ knots are negative. They did not report the sign for the trefoils. The trefoils and the 5₁ knots are torus knots made by the same reaction, so we assumed that the trefoil crossings are also positive. TopoStats calls 76 of the 98 labeled trefoil crossings positive (Table 2), which supports this assumption, although the labels were not derived from these calls.

**Table 2.**
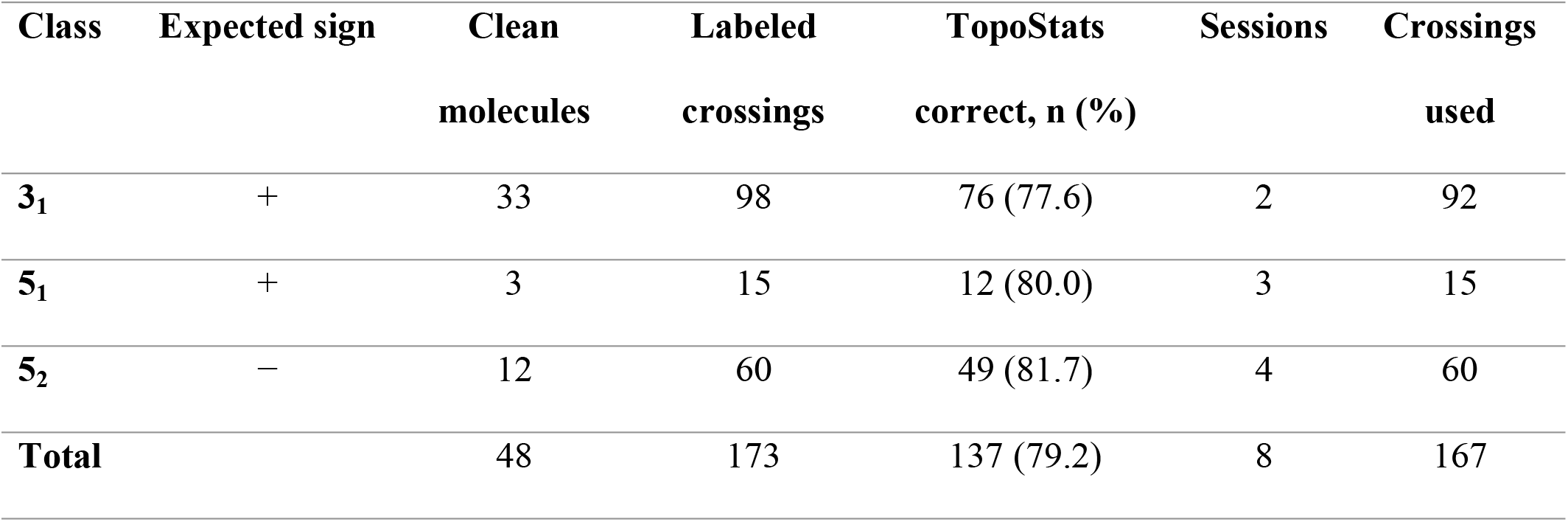
Crossing labels obtained from knot handedness.

| **Class** | **Expected sign** | **Clean molecules** | **Labeled crossings** | **TopoStats correct, n (%)** | **Sessions** | **Crossings used** |
|---|---|---|---|---|---|---|
| **3<sub>1</sub>** | + | 33 | 98 | 76 (77.6) | 2 | 92 |
| **5<sub>1</sub>** | + | 3 | 15 | 12 (80.0) | 3 | 15 |
| **5<sub>2</sub>** | − | 12 | 60 | 49 (81.7) | 4 | 60 |
| **Total** |  | 48 | 173 | 137 (79.2) | 8 | 167 |

The labels apply only to clean molecules, in which each crossing region holds exactly one crossing. A molecule was called clean when TopoStats found as many crossing regions as the minimal crossing number of its sample (3 for 3₁, and 5 for 5₁ and 5₂), when every region contained exactly two strands, and when the molecule was among the 721 molecules of Table 1. In a clean molecule, the expected sign *s* of every crossing is known. TopoStats reports its own sign *ŝ* for each crossing, computed from the directions of its traced strands and from its choice of upper strand. If *ŝ* = *s*, TopoStats placed the correct strand on top. If *ŝ* ≠ *s*, the other strand lies on top. Crossings without a reported sign were not used.

This procedure gave 48 clean molecules (33 trefoils, 3 torus knots and 12 twist knots) with 173 labeled crossings (Table 2). In 22 molecules, every TopoStats sign agreed with the expected sign, and in 24 molecules some signs disagreed. In the remaining two molecules, both trefoils, every sign disagreed. Such a molecule is either a mirror-image knot, which the sample should not contain, or the result of a systematic tracing failure, so its labels cannot be trusted. These two molecules were excluded from the learning experiments, which therefore used 167 crossings from 46 molecules and eight imaging sessions.

#### Learned crossing classifier

To test whether a learned model can do better than the width rule, a small convolutional network was trained to decide which of two strands lies on top at a crossing. The width rule measures the width of each strand as the full width at half maximum (FWHM) of its height profile across the crossing [8]. The network instead received an image patch of 64 × 64 nm centered on the crossing region, with three channels. The first channel held the height image, and the other two marked the paths of the two strands (S2 Fig).

The network was trained in three ways. In the first, it learned only from the 167 crossings whose upper strand was known from knot handedness (see the previous section). In the second, it learned from the decisions that TopoStats makes at every crossing, which are far more numerous but correct for only 79% of the 173 labeled crossings (Table 2). In the third, it learned first from the TopoStats decisions and then from the 167 known crossings. This combines many noisy labels with a few exact ones, a common strategy when exact labels are scarce [34].

The crossings were divided into five folds by imaging session with GroupKFold, so all crossings of one session fell in the same fold. Every crossing was therefore predicted by a model that had seen no crossing of its session, neither as a known crossing nor as a TopoStats decision.

TopoStats measures widths on the original images, whose pixels are 0.29–0.98 nm, whereas the network saw patches resampled to 1 nm per pixel. Patches were therefore also sampled at 0.5 nm per pixel, to test whether the coarser sampling removes useful detail. The architecture, training settings and sample counts are given in S2 Appendix.

#### Knot type and crossing counts

To test whether better connections give correct knot types, the knot type was computed with energy-based pairing in place of the pairing of TopoStats. The other two steps followed the same ideas as TopoStats. First, the upper strand at each crossing was decided by the width rule.

Because the strands now follow the new connections, their widths were measured again along these connections. Second, the knot type was computed with Topoly [16], the tool that TopoStats also uses. For this, the molecule was turned into a three-dimensional curve, in which the strands in each crossing region are lifted to different heights in the order set by the width rule. This records which strand passes over which, which a flat trace cannot do. Topoly then computed the HOMFLY-PT polynomial of the curve, which identifies the knot or link type [9]. Mirror images were not distinguished, because the evaluation concerns the knot type and not its handedness.

Molecules with more than 14 crossings were not evaluated, and each Topoly calculation was stopped after 20 s, because larger tangles can make the computation run indefinitely. Both cases counted as errors, as did molecules without a valid pairing. A molecule was counted as correct when its computed knot type equaled the type of its sample. TopoStats was scored on the same molecules by the type it reports.

To find where the pipeline fails, the crossings between strands inside the crossing regions were counted for each molecule and compared with the minimal crossing number of its type (3 for 3₁, 4 for 4²₁, and 5 for 5₁ and 5₂). A molecule with fewer crossings than this number has lost crossings before any upper strand was chosen, so it cannot receive its correct type, however well the crossings are ordered. A molecule with exactly the minimal number *n* of crossings receives its correct type only if every crossing is ordered correctly, or if every crossing is reversed, which gives the mirror image and therefore the same type. If each crossing is ordered correctly with probability *p*, independently of the others, the type is correct with a probability of about *p*ⁿ + (1 − *p*)ⁿ [8]. Here *p* was taken as the accuracy of the width rule on the labeled crossings. This expected share was compared with the observed share of correct types. An observed share below the expected one would indicate errors other than the ordering of crossings.

### Comparison with TopoStats and statistical analysis

TopoStats was always evaluated on the same molecules or crossings as the method it was compared with, so both were judged on identical cases. Its knot types and crossing decisions were read from the .topostats file of each image. A TopoStats type was counted as correct only when it matched the true type exactly, including the number of loops. For example, a single unknotted ring that TopoStats reported as two separate unknotted rings was counted as misclassified, because the trace had split one molecule into two. A molecule to which TopoStats assigned no knot type was also counted as misclassified.

The uncertainty of each result is given as a 95% confidence interval (CI), estimated by bootstrap resampling [35]. The data were resampled with replacement 2,000 times, the result was recomputed each time, and the interval was taken between the 2.5th and 97.5th percentiles of these values. Molecules imaged in the same session are not independent, because they share the conditions of that session. For whole-molecule classification, whole sessions were therefore resampled instead of single molecules. For each class, the sessions containing it were drawn with replacement, and all molecules of that class from the drawn sessions were used, so that every class was present in every resample. For crossings, and for the pairing and knot-type evaluations, single crossings or molecules were resampled.

Two methods evaluated on the same items were compared with the exact McNemar test [36]. The test uses only the discordant items, which one method classifies correctly and the other does not, and asks whether one method gets such items right more often than chance would explain. For whole molecules, the compared quantity was the knots-only accuracy, the share of the 293 knot molecules assigned their correct class. Because many comparisons were made, *p* values were adjusted for multiple testing with the Holm procedure [37] within each family of related comparisons. The four families were the 36 knots-only comparisons of the whole-molecule classifiers, including the control analyses, the 15 comparisons of the crossing classifiers, the seven comparisons of the pairing evaluation and the eight comparisons of the knot-type evaluation. A difference was considered significant when its adjusted *p* value was below 0.05.

Comparisons within subgroups of crossings were exploratory and are reported without adjustment. No missing values were imputed, because every failure was counted as an error. Analyses were run in Python 3.14.8 with PyTorch 2.14.1, timm 1.0.30, scikit-learn 1.9.1, h5py 3.16.0 and Topoly 1.1.0. All analysis code is available in the repository given in the Data Availability Statement.

### Use of AI tools

The authors used Claude (Anthropic), a large language model, in two ways. First, it assisted in writing the code that draws the illustrative figures, namely the knot diagrams (Fig 1), the pipeline diagram (Fig 2) and the pairing schematic (Fig 3). Second, it assisted in writing the analysis code. The authors checked all such code by running it on the data and comparing its outputs with the expected counts and with the released TopoStats output. The authors reviewed all AI-assisted code and figures and take full responsibility for the content of the article.

## Results

### Accuracy of the existing pipeline

Before the new methods were tested, the accuracy of TopoStats was measured on the 721 molecules of Table 1, to set the baseline for all later comparisons. TopoStats assigned the correct type to only 44 of the 588 knots and catenanes (7.5%), but to 115 of the 133 unknots (86.5%, Table 1). It therefore identified simple rings reliably but failed on most knotted and linked molecules.

### The session-aware protocol exposes session leakage

The session-aware protocol was first applied to the three whole-molecule classifiers, and Table 3 compares their results under the image split and the session split. Under the image split, the classifiers reached knots-only balanced accuracies of 57.7–67.4%. Under the session split, the same models reached only 27.0–32.0%, close to the chance level of 25%. The image split therefore overestimated performance on the same molecules by 40.3 percentage points for ResNet-18 (95% CI 23.7–44.1), by 32.2 for ConvNeXt-Tiny (3.2–38.8) and by 30.2 for DINOv2 (6.6–35.2). These intervals were computed by resampling imaging sessions, and none of them includes zero, so the gap is not a chance result. The knots-only macro AUC showed the same pattern. It was 0.86–0.89 under the image split but only 0.49–0.57 under the session split, close to 0.5. On new sessions, therefore, not only the predicted class but also the predicted probabilities carried almost no information about knot type.

**Table 3.** Whole-molecule classification under the image split and the session split.

| **Model** | **Split** | **Accuracy (%)** | **Balanced accuracy (%)** | **Macro-F1 (%)** | **Knots-only balanced accuracy (%)** | **Knots-only macro AUC** |
|---|---|---|---|---|---|---|
| **ResNet-18** | image | 89.2 | 73.8 | 73.2 | 67.4 [53.7–75.9] | 0.89 |
| **ConvNeXt-Tiny** | image | 87.9 | 71.2 | 71.2 | 64.2 [45.3–73.2] | 0.89 |
| **DINOv2 + LR** | image | 85.2 | 65.9 | 65.9 | 57.7 [41.7–61.8] | 0.86 |
| **ResNet-18** | session | 75.2 | 41.3 | 41.5 | 27.0 [20.7–40.9] | 0.49 |
| **ConvNeXt-Tiny** | session | 77.5 | 45.5 | 45.6 | 32.0 [23.2–53.7] | 0.53 |
| **DINOv2 + LR** | session | 75.5 | 41.7 | 41.8 | 27.5 [22.9–39.7] | 0.57 |
| **TopoStats** | – | 22.1 | 29.9 | 22.0 | 36.4 [23.9–42.9] | – |

Overall accuracy concealed this failure. Under the session split, it remained at 75.2–77.5%, because catenanes make up 428 of the 721 molecules (59.4%) and each model recognized them in 98.4–99.5% of cases under either split (S1 Table). The main result therefore came from the four knot types that cannot be separated by size. On these types, TopoStats reached a knots-only balanced accuracy of 36.4% (95% CI 23.9–42.9) and classified 142 of the 293 knot molecules (48.5%) correctly. No classifier tested under the session split did better. ConvNeXt-Tiny classified 45.7% of the knot molecules correctly and did not differ significantly from TopoStats (*p* = 0.40). ResNet-18 (41.3%) and DINOv2 (42.0%) were less accurate than TopoStats before adjustment for multiple comparisons (*p* = 0.010 and 0.030), but not after it (adjusted *p* = 0.26 and 0.62). Under the image split, by contrast, all three classifiers were more accurate than TopoStats (65.5–74.1%, adjusted *p* < 0.001). A study that reported only the image split would therefore have concluded that learned classifiers outperform the existing pipeline.

The errors under the session split followed the structure of the sessions (Fig 4). None of the three models recognized any trefoil. Because 38 of the 41 trefoils belong to one session, the training folds contain only three trefoils whenever that session is tested (S3 Table). The two five-crossing knots were systematically confused with each other. Under the session split, ResNet-18 assigned 41 of the 63 torus knots to 5₂ and 29 of the 56 twist knots to 5₁, compared with 13 and 12 under the image split. These errors were concentrated in the two largest sessions of five-crossing knots, the torus knots of 21 August 2023 and the twist knots of 17 August 2023. All three models classified these two sessions with 0–5% accuracy (S2 Table). Apart from three twist knots, these two sessions hold the only five-crossing knots imaged in 2023, and they fall in different folds.

**Fig 4.**
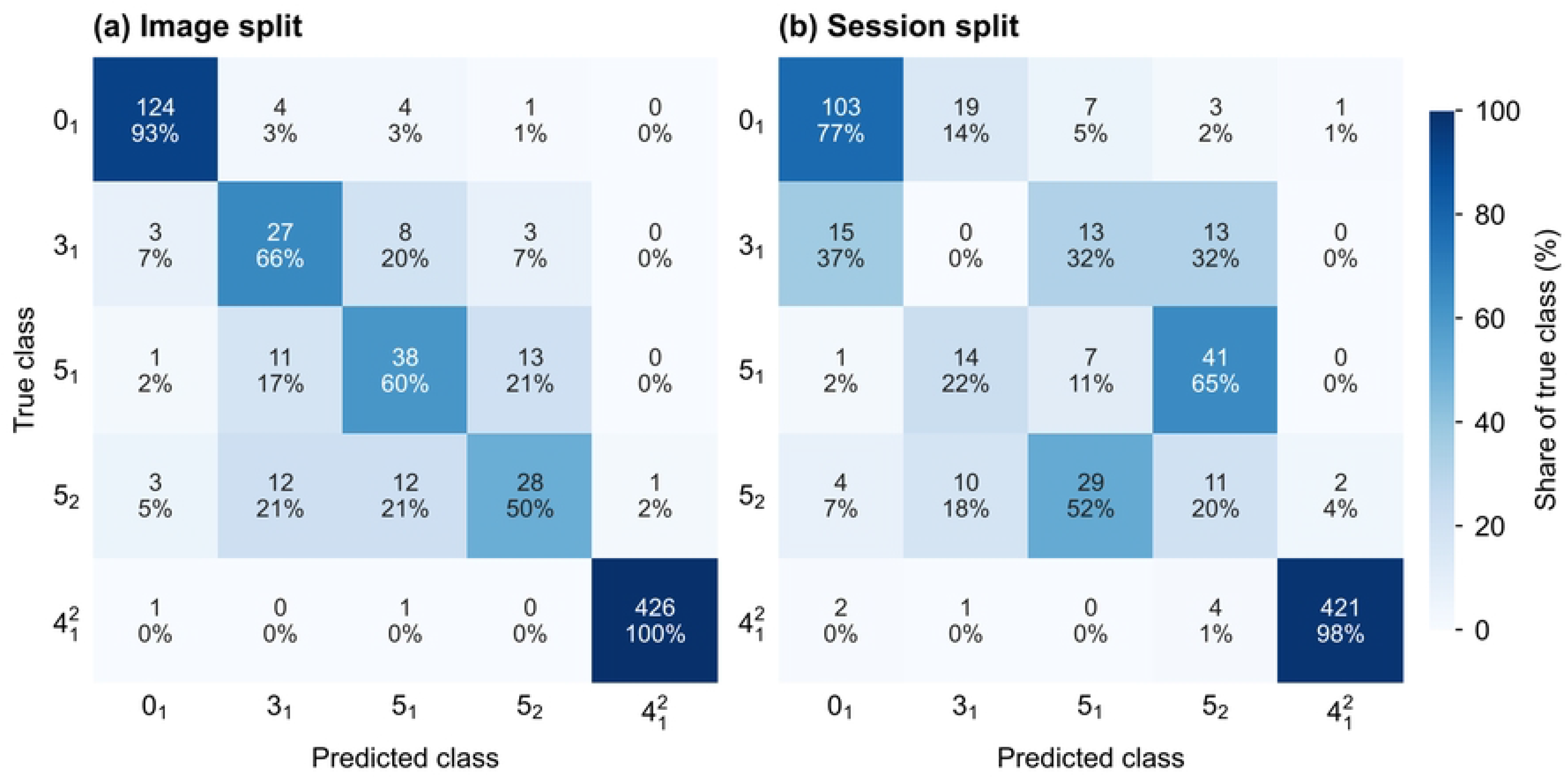
Confusion matrices of ResNet-18 under the two splits. (a) Image split. (b) Session split. Rows are true classes and columns are predicted classes. Each cell gives the number of molecules and the row percentage.

Whenever one of them was tested, the five-crossing knots from 2023 in the training data therefore belonged almost entirely to the other type. The models assigned the type that matched the imaging year, not the knot.

The session probe confirmed that images carry a signature of their session (Table 4). Trained on images of a single preparation, the probe identified the imaging session above chance in all eight tests (*p* ≤ 0.025). This also held in the five tests whose sessions all used the same imaging buffer. For nicked catenanes imaged in the Mg^2+^/Ni^2+^ buffer, the probe reached 58.2% against a chance level of 33.3%, and for nicked unknots 94.7% against 50%. Molecules of one preparation share their topology and supercoiling, so the probe recognized the session without any information about the knot.

**Table 4.** Session probe.

| **Preparation** | **Buffer** | **Molecules** | **Sessions** | **Balanced accuracy (%)** | **Chance (%)** | ***p*** |
|---|---|---|---|---|---|---|
| **Nicked catenane** | both | 239 | 5 | 42.5 | 20.0 | 0.005 |
| **Nicked catenane** | Mg<sup>2+</sup>/Ni<sup>2+</sup> | 201 | 3 | 58.2 | 33.3 | 0.005 |
| **Nicked catenane** | Ni<sup>2+</sup> | 38 | 2 | 75.3 | 50.0 | 0.025 |
| **Supercoiled catenane** | Mg<sup>2+</sup>/Ni<sup>2+</sup> | 176 | 2 | 74.2 | 50.0 | 0.005 |
| **Torus knot (5<sub>1</sub>)** | both | 59 | 3 | 54.9 | 33.3 | 0.010 |
| **Twist knot (5<sub>2</sub>)** | both | 53 | 3 | 64.6 | 33.3 | 0.005 |
| **Nicked unknot** | Mg<sup>2+</sup>/Ni<sup>2+</sup> | 64 | 2 | 94.7 | 50.0 | 0.005 |
| **Supercoiled unknot** | Mg<sup>2+</sup>/Ni<sup>2+</sup> | 69 | 2 | 76.0 | 50.0 | 0.005 |

The control analyses tested whether a different training setting could raise performance on new sessions (S1 Appendix). Flip augmentation, training for 15 or 60 epochs, normalized or binary input images and sampling at 0.5 nm per pixel all left the knots-only balanced accuracy between 25.1% and 36.1%. After adjustment for multiple comparisons, none of these runs differed significantly from the reference run of the same model. The highest value, 36.1% for ConvNeXt-Tiny trained for 15 epochs, only equaled that of TopoStats (36.4%).

### Energy-based pairing corrects most loop errors

Energy-based pairing was compared with TopoStats on how often each traced the loops expected for the sample (Table 5). Energy-based pairing traced the correct loops for 614 of the 721 molecules (85.2%, 95% CI 82.5–87.9), and TopoStats for 311 (43.1%, 39.5–46.7). For knots and catenanes together, the share was 493 of 588 (83.8%) with energy-based pairing and 186 (31.6%) with TopoStats (adjusted *p* < 0.001). The gain came from the types that TopoStats traces least accurately. For catenanes, the share was 85.0% with energy-based pairing and 19.6% with TopoStats, and for torus knots 77.8% and 44.4% (adjusted *p* < 0.001 and *p* = 0.001). For twist knots, the share was also higher with energy-based pairing (82.1% vs. 67.9%), but the difference was not significant after adjustment (adjusted *p* = 0.23). For trefoils and unknots, TopoStats was slightly more accurate, but the two methods did not differ significantly (87.8% vs. 82.9% and 94.0% vs. 91.0%, adjusted *p* = 0.85 for both). Energy-based pairing found no valid solution for 101 molecules (14.0%), and these molecules were counted as errors. In 58 of them, the skeleton contained an open end or a detached fragment, and in 43, a crossing region had an odd number of segment ends.

**Table 5.** Molecules whose traced loops have the expected number and lengths.

| **Class** | ***n*** | **TopoStats (%)** | **Energy-based pairing (%)** | **Only pairing right** | **Only TopoStats right** | **Adjusted *p*** |
|---|---|---|---|---|---|---|
| **0<sub>1</sub>** | 133 | 94.0 [90.2–97.7] | 91.0 [85.7–95.5] | 5 | 9 | 0.85 |
| **3<sub>1</sub>** | 41 | 87.8 [78.1–97.6] | 82.9 [70.7–92.7] | 1 | 3 | 0.85 |
| **5<sub>1</sub>** | 63 | 44.4 [31.7–57.1] | 77.8 [66.7–87.3] | 27 | 6 | 0.001 |
| **5<sub>2</sub>** | 56 | 67.9 [55.4–80.4] | 82.1 [71.4–91.1] | 12 | 4 | 0.23 |
| **4<sup>2</sup><sub>1</sub>** | 428 | 19.6 [16.1–23.6] | 85.0 [81.5–88.3] | 289 | 9 | < 0.001 |
| **Knots and catenanes** | 588 | 31.6 [28.1–35.5] | 83.8 [80.8–86.7] | 329 | 22 | < 0.001 |
| **All** | 721 | 43.1 [39.5–46.7] | 85.2 [82.5–87.9] | 334 | 31 | < 0.001 |

### Crossing resolution limits the knot type

#### Accuracy of crossing order

The labels from knot handedness first gave the accuracy of the width rule (Table 2). On the 173 labeled crossings, TopoStats placed the correct strand on top in 137 cases (79.2%). This is close to the 82% that Holmes et al. [8] measured against hand labels, and the accuracy was similar for the three knots (77.6–81.7%). TopoStats also reports a reliability score for each decision [8], and this score separated accurate from inaccurate decisions. Below the threshold of 0.263 suggested by Holmes et al. [8], TopoStats was correct for 86 of 115 crossings (74.8%), and above it for 51 of 58 (87.9%). On the 167 crossings used for learning, which exclude the two trefoils whose signs all disagreed, TopoStats was correct for 82.0% (95% CI 76.0–87.4).

The learned crossing classifier did not do better than the width rule (Table 6). Trained only on the 167 known crossings, the network performed at chance level (49.7% at 1 nm and 50.9% at 0.5 nm). Trained on the decisions of TopoStats, it was correct for 77.2% of the crossings at either resolution. Further training on the known crossings gave 78.4% at 1 nm and 77.2% at 0.5 nm.

**Table 6.** Crossing classifiers and TopoStats on the same 167 labeled crossings.

| **Resolution** | **Training** | **Accuracy (%)** | **Same decision as TopoStats (%)** | **Only network right** | **Only TopoStats right** | **Adjusted *p*** |
|---|---|---|---|---|---|---|
| **1 nm** | known crossings only | 49.7 [42.5–57.5] | 49.7 | 15 | 69 | < 0.001 |
|  | TopoStats decisions | 77.2 [70.7–83.2] | 83.2 | 10 | 18 | 1.00 |
|  | TopoStats decisions, then known crossings | 78.4 [71.9–84.4] | 84.4 | 10 | 16 | 1.00 |
| **0.5 nm** | known crossings only | 50.9 [43.7–58.1] | 49.7 | 16 | 68 | < 0.001 |
|  | TopoStats decisions | 77.2 [70.7–83.2] | 85.6 | 8 | 16 | 1.00 |
|  | TopoStats decisions, then known crossings | 77.2 [70.7–83.2] | 82.0 | 11 | 19 | 1.00 |
| **–** | TopoStats | 82.0 [76.0–87.4] | – | – | – | – |

After adjustment for multiple comparisons, none of these four networks differed significantly from TopoStats (adjusted *p* = 1.00 for each). The networks trained on the TopoStats decisions made the same decision as TopoStats on 82.0–85.6% of the crossings. They matched TopoStats on crossings that it rated as reliable, but they were less accurate on crossings that it rated as unreliable (S2 Appendix). The networks therefore learned the width rule together with its errors, and they did not correct the crossings on which the rule fails.

#### Knot type with energy-based pairing

Correctly traced loops did not yield the correct knot type. With energy-based pairing and the width rule, the type was correct for 36 of 588 knots and catenanes (6.1%, 95% CI 4.3–8.2), against 44 (7.5%, 5.4–9.7) for TopoStats on the same molecules (Table 7, adjusted *p* = 1.00). No class differed between the two methods after adjustment. Of the 493 knots and catenanes whose loops the pairing traced correctly, only 36 (7.3%) received the correct type. Every unknot for which the pairing found a solution was typed as an unknot, so the pipeline did not introduce crossings that change the type of a simple molecule. The incorrect types were simpler than the true ones (S4 Table). Of the 428 catenanes, 165 were typed as the Hopf link 2^2^ and 101 as two unlinked circles, and five-crossing knots were most often typed as unknots (58 of 119).

**Table 7.** Molecules assigned the type of their sample.

| **Class** | ***n*** | **TopoStats (%)** | **Energy-based pairing and width rule (%)** | **Only new method right** | **Only TopoStats right** | **Adjusted *p*** |
|---|---|---|---|---|---|---|
| **0<sub>1</sub>** | 133 | 86.5 [80.5–91.7] | 91.0 [85.7–95.5] | 7 | 1 | 0.51 |
| **3<sub>1</sub>** | 41 | 43.9 [29.3–58.5] | 29.3 [14.6–43.9] | 2 | 8 | 0.66 |
| **5<sub>1</sub>** | 63 | 6.3 [1.6–12.7] | 3.2 [0.0–7.9] | 1 | 3 | 1.00 |
| **5<sub>2</sub>** | 56 | 8.9 [1.8–16.1] | 7.1 [1.8–14.3] | 2 | 3 | 1.00 |
| **4<sup>2</sup><sub>1</sub>** | 428 | 4.0 [2.3–5.8] | 4.2 [2.6–6.1] | 9 | 8 | 1.00 |
| **Knots (3<sub>1</sub>, 5<sub>1</sub>, 5<sub>2</sub>)** | 160 | 16.9 [11.3–23.1] | 11.3 [6.3–16.3] | 5 | 14 | 0.51 |
| **Knots and catenanes** | 588 | 7.5 [5.4–9.7] | 6.1 [4.3–8.2] | 14 | 22 | 1.00 |
| **All** | 721 | 22.1 [18.9–25.2] | 21.8 [18.7–24.8] | 21 | 23 | 1.00 |

#### Lost crossings

Counting the crossings in each molecule showed where the pipeline fails (Table 8). These counts were made for the molecules for which energy-based pairing found a solution. Fewer crossings than the minimal crossing number were found in 308 of 368 catenanes (83.7%), 32 of 50 torus knots (64.0%) and 20 of 46 twist knots (43.5%), but in none of the 35 trefoils. Across knots and catenanes, 360 of 499 molecules (72.1%) had lost crossings before any upper strand was chosen. None of these molecules can receive its correct type, and none did. The crossings are lost because TopoStats merges crossing points that lie within 7 nm of each other into a single crossing region [8]. When two strands cross twice in close succession, both crossings fall into one region. That region then has only four segment ends, the same as a single crossing, so at most one of the two crossings can be recovered.

**Table 8.** Crossings found in each molecule compared with the minimal crossing number n.

| **Class** | **Molecules** | **No valid pairing** | **Fewer than *n*** | **Exactly *n*** | **More than *n*** | **Correct type with exactly *n*, n (%)** | ***p<sup>n</sup>* + (1 − *p*)<sup>*n*</sup>** |
|---|---|---|---|---|---|---|---|
| **3<sub>1</sub> (*n* = 3)** | 41 | 6 | 0 | 31 | 4 | 11 (35.5) | 0.56 |
| **4<sup>2</sup><sub>1</sub> (*n* = 4)** | 428 | 60 | 308 | 42 | 18 | 12 (28.6) | 0.45 |
| **5<sub>1</sub> (*n* = 5)** | 63 | 13 | 32 | 7 | 11 | 1 (14.3) | 0.37 |
| **5<sub>2</sub> (*n* = 5)** | 56 | 10 | 20 | 14 | 12 | 1 (7.1) | 0.37 |

Where the minimal number of crossings was found, errors in ordering individual crossings set the next limit. The width rule ordered 82% of the labeled crossings correctly (*p* = 0.82). At this accuracy, the probability that all crossings of a molecule are ordered consistently is 0.56 for three crossings, 0.45 for four and 0.37 for five. The observed shares of correct types were lower in every class (Table 8). They were 11 of 31 trefoils (35.5%), 12 of 42 catenanes (28.6%), 1 of 7 torus knots (14.3%) and 1 of 14 twist knots (7.1%). This points to further errors, either in the pairing or in ordering crossings along the new connections. The accuracy of ordering along the new connections could not be measured, because the available labels refer to the strand paths traced by TopoStats.

## Discussion

### The session-aware protocol prevents an overstated conclusion

The session-aware protocol changed the conclusion of the classifier evaluation. Under the image split, all three classifiers appeared to read the knot type from AFM images, and their knots-only balanced accuracy exceeded that of TopoStats by 21–31 percentage points. Under the session split, none of them outperformed TopoStats, and all were close to chance on knots. What remained was the recognition of catenanes, which differ from the other molecules in size, and a partial recognition of unknots. Without the knots-only metric, this failure would have been hidden by an overall accuracy above 75%. The gap between the two splits was of similar size for two convolutional networks and a frozen self-supervised transformer. It therefore arises from the data and the evaluation, not from a particular architecture.

The most likely cause is the way AFM data are acquired. Each sample is imaged in few sessions, and sessions differ in tip condition, surface preparation and imaging buffer. Under the image split, molecules from the same session appear in both the training and the test folds. Any image feature that identifies the session then also identifies the knot type, and a classifier can learn this shortcut [21]. The same effect has been found in chest radiographs, where classifiers learned features of the hospital that took the image [24]. Three observations support this explanation.

First, the session probe showed that such features exist, even among sessions that used the same buffer. Second, the models confused the two five-crossing knots in a pattern that followed the imaging dates of the training sessions, whereas a model without useful information would have guessed between them at random. Third, none of the control analyses raised session-split performance above that of TopoStats, so the failure is not explained by the training settings.

These results do not show that the knot type cannot be learned from AFM images. They show that this dataset cannot establish whether it can. The trefoils come almost entirely from one session, so no session-split test can evaluate them. The five-crossing knots span four sessions each, which is too few for a classifier to learn features that stay the same across sessions. The protocol itself is not specific to knots. It applies to any AFM study in which the sample and the imaging session are confounded, and the session probe can test for this confounding before any classifier is trained.

### Energy-based pairing as a replacement for direction-based pairing

Energy-based pairing raised the share of knots and catenanes with correctly traced loops from 32% with TopoStats to 84%. The gain was largest for catenanes and torus knots, the types that TopoStats traces least accurately, and the method was not significantly worse than TopoStats for any type. The method needs no training data, and it does not need to know which sample a molecule comes from. It uses only the expected lengths of the molecules in a dataset and the stiffness of DNA. Both are known in advance in most AFM studies of plasmids and their recombination products, so only the expected lengths need to be set for a new dataset. The main failure of the method was a skeleton with an open end, a detached fragment or a crossing region with an odd number of segment ends. This affected 14% of molecules and might be reduced by repairing the skeleton before pairing.

Correct loops, however, are necessary but not sufficient for the correct knot type. On its own, the pairing did not raise the share of correct knot types. It is therefore most useful where the number and length of loops are the quantities of interest, for example when catenated products are counted.

### Crossing detection, not crossing order, limits the pipeline

The measured limit locates the remaining failure. Of the knots and catenanes for which the pairing found a solution, 72% had fewer crossings than their type requires before any crossing was ordered, and none of them received the correct type. Most catenanes and torus knots were lost at this step, because closely spaced crossings merged into regions from which not all crossings can be recovered. Errors in ordering crossings matter only for the molecules whose crossings were all found. Raising the per-crossing accuracy from 82% to 90% would raise the probability of a consistent order from 0.56 to 0.73 for three crossings and from 0.37 to 0.59 for five. This would help only the 94 knots and catenanes whose crossings were all found, 19% of those with a solution, so it would raise the share of correct types by at most a few percentage points.

The crossing classifier suggests that even this smaller gain is hard to obtain by learning. Trained on the few known crossings, the network learned nothing that transferred to new sessions. Trained on the decisions of the width rule, it reproduced the rule together with its errors. The known crossings are also the isolated ones, on which the rule is already right about four times in five. The largest gain is therefore available from resolving dense crossing regions into individual crossings. The heights measured inside a region and the bending energy of the pairing method are natural inputs for this step. Labels from knot handedness could be used to test such a method wherever the topology and chirality of a sample are known. The 167 known crossings of this study already provide a benchmark on which any method for crossing order can be compared with the width rule.

### Recommendations

Four practices would make machine-learning results on AFM images easier to trust and to compare. First, models should be tested on imaging sessions not used for training, and papers should report how many sessions each type spans. Second, results should include a metric that excludes types that can be separated by size or by another simple cue. Third, learned models should be compared with the existing rule on the same molecules, with a paired test. Fourth, pipelines should be evaluated step by step, so that each gain or loss can be traced to the step that causes it. In addition, datasets for training topology classifiers should image each type over many sessions and, where possible, several types in each session, so that session and type are no longer confounded.

### Limitations and future work

The dataset limits what this study can show. It contains few imaging sessions per type and comes from one laboratory and one family of instruments. The session split therefore cannot fully separate shortcut learning from the effect of too few sessions, and the size of the session effect may differ in other laboratories. Because only 12 sessions contain knots, the confidence intervals of the session split are wide, and they do not exclude knots-only balanced accuracies of up to about 54%. The McNemar tests treat molecules and crossings as independent, so their *p* values may be too small where items from the same session are correlated. Datasets with many sessions per type, ideally from several laboratories, would allow these effects to be measured more precisely.

The crossing labels have four limits. First, they cover only clean molecules, in which the crossings are well separated, so the measured accuracies of crossing order are probably upper bounds for crossings in dense regions. Second, the sign of the trefoil crossings was inferred from the reported chirality of the 5₁ torus knot, not measured on the trefoil sample. If the trefoils had the opposite chirality, their labels would be inverted, while the labels of 5₁ and 5₂ would not change. Third, the labels depend on the strand directions of the TopoStats trace. Fourth, 89 of the 167 known crossings come from one trefoil session. In the pairing evaluation, the loop test shares its length targets with the energy and therefore favors the new method, but the knot-type test does not. Finally, all accuracies assume that every molecule in a sample has the type of that sample. The recombination products were purified but are not guaranteed to be pure, so some apparent errors may be correct decisions on impure molecules. TopoStats identified 86.5% of the unknot controls correctly, so at most 13.5% of the unknot molecules can be impurities.

## Conclusions

This study presented three methods for evaluating and improving the automated identification of DNA knot types in AFM images. The session-aware test protocol showed that whole-molecule classifiers learn the imaging session rather than the knot type, and that tests which divide the data by image overstate their performance. Energy-based pairing corrected most loop errors without any training data. The measured crossing-resolution limit showed that the pipeline is limited mainly by the merging of closely spaced crossings rather than by errors in their order. The protocol, the crossing labels and the code are openly available, so that future methods can be compared with these baselines and directed to the detection of dense crossings.

## Acknowledgments

We thank the authors of the public dataset for releasing the AFM images and the TopoStats output openly, which made this study possible.

## Supporting information

**S1 Appendix. Control analyses for the whole-molecule classifiers.** Methods and results of the control analyses of flip augmentation, training length, input encoding and sampling density, including the table of knots-only metrics and McNemar tests.

**S2 Appendix. Details of the crossing classifier.** Sample extraction, network architecture, training settings, the three training regimes and results by knot type, reliability score and imaging session.

**S1 Fig.**
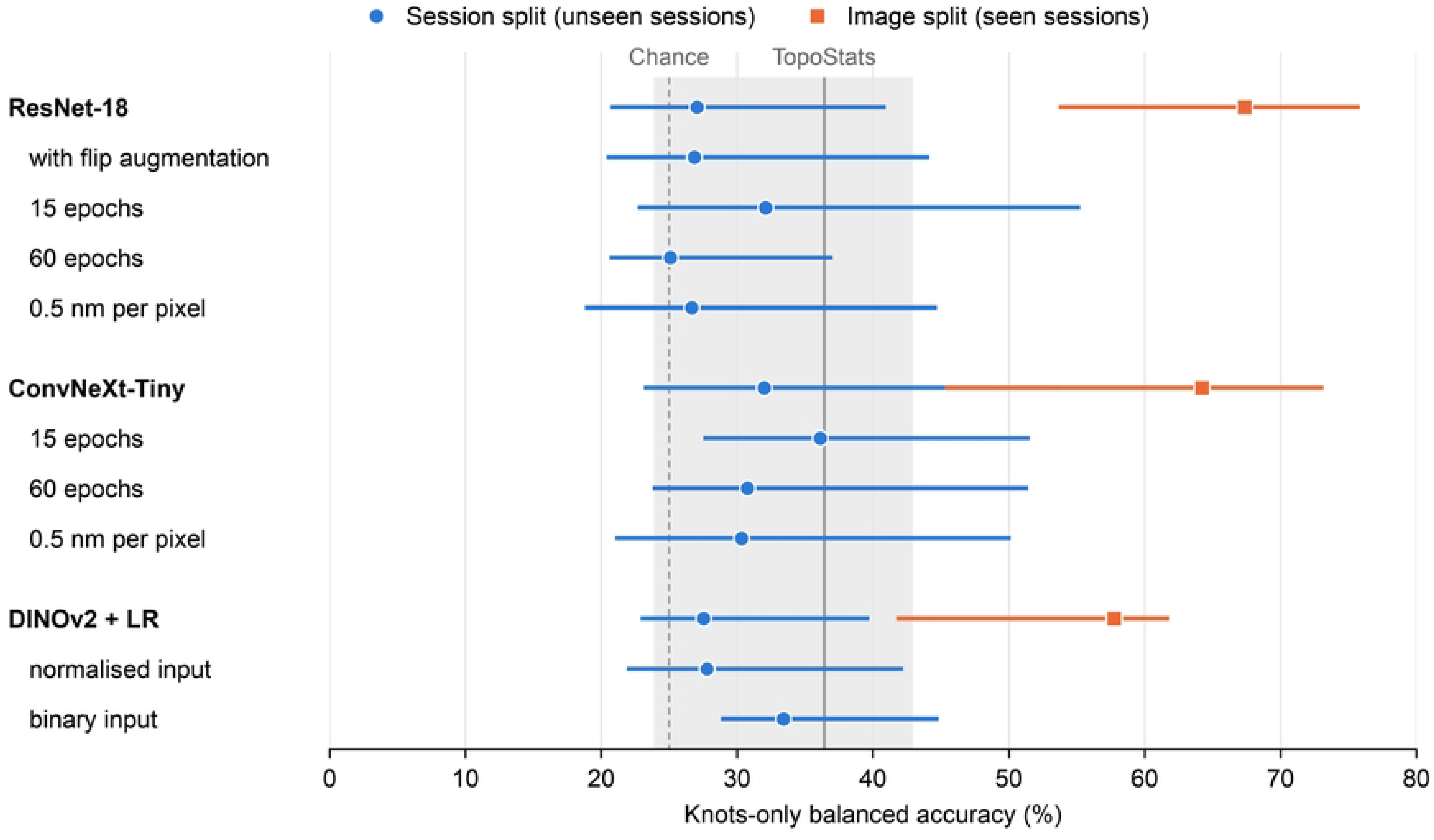
Knots-only balanced accuracy of all whole-molecule classifier runs. Image split (orange squares) and session split (blue circles), including the control runs. Error bars show 95% confidence intervals from resampling imaging sessions. The dashed line marks chance (25%) and the solid grey line TopoStats on the same molecules, with its confidence interval shaded.

**S2 Fig.**
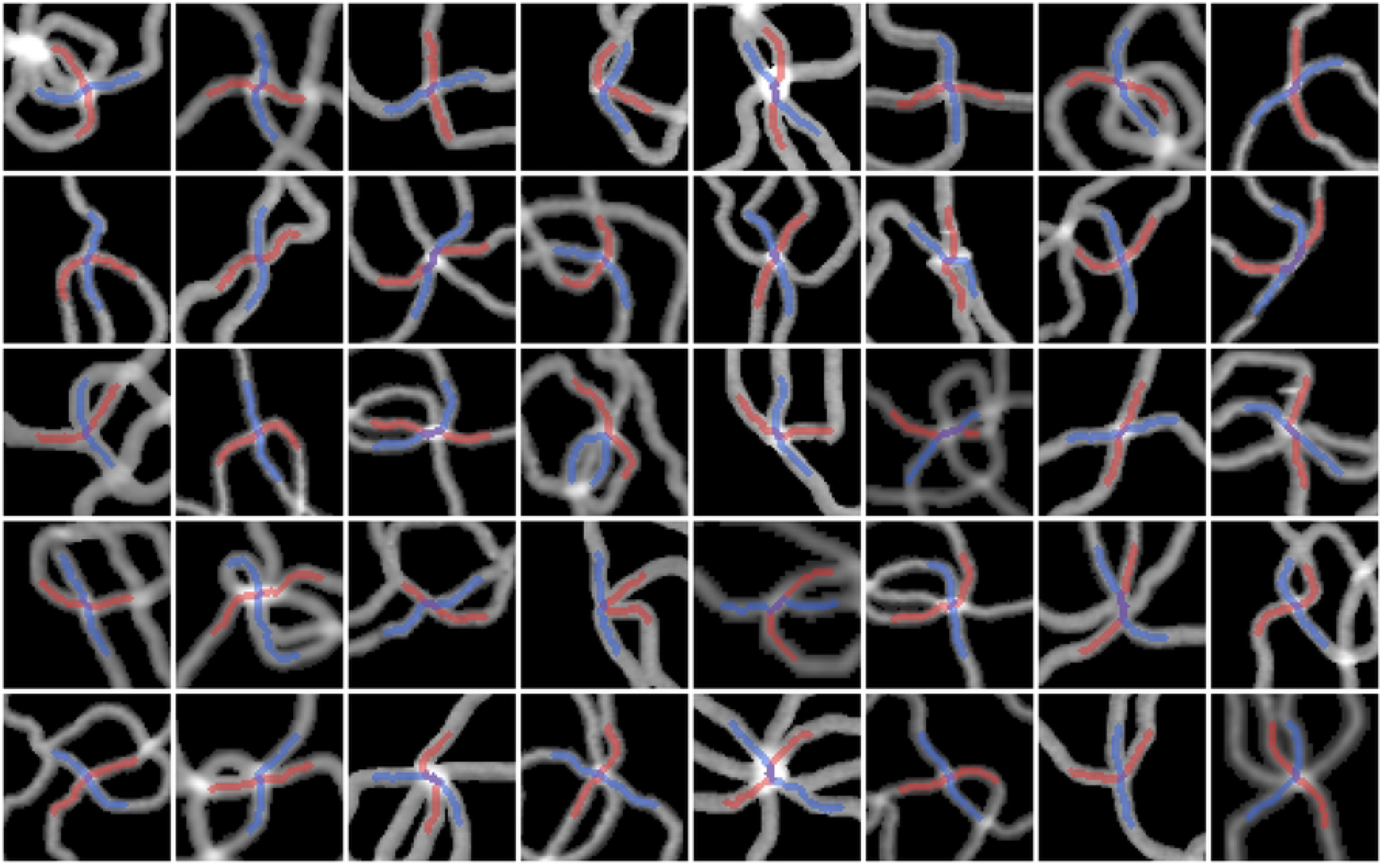
Examples of crossing patches. Forty randomly chosen crossing patches at 1 nm per pixel, each covering 64 × 64 nm. Grey shows the height image on a linear scale from −1 to 5 nm, red the path of strand A and blue the path of strand B.

**S1 Table. Recall per class under the image split and the session split.**

**S2 Table. Accuracy per imaging session under the session split.** 1 nm height input.

**S3 Table. Composition of the five session folds.** Number of molecules per class in each fold.

**S4 Table. Types assigned by energy-based pairing and the width rule.** Per class, on the 721 molecules.

